# Replicative and chronological epigenetic age track genome instability in opposite directions across cancers

**DOI:** 10.64898/2026.07.29.741419

**Authors:** Aditri Mukherji, Subhayan Chattopadhyay

## Abstract

Mitotic epigenetic clocks are increasingly applied to tumors, but what a tumor’s advance over its own normal tissue records, and whether this differs from what chronological clocks record, has been examined in tumor samples alone. We compared four mitotic clocks with Horvath’s pan tissue clock in 506 matched tumor and normal pairs from 13 adult solid cancers and in 1,731 pediatric tumors, and partitioned the tumor minus normal difference into proliferation, cell composition, global methylation change and genome instability. Mitotic age was higher in the tumor in every adult cancer type, robust to purity and composition. Global methylation change and composition accounted for most of this advance (partial R² up to 25% and 18%), whereas tumor proliferation contributed 2 to 7% and generalised across cancer types only for stemTOC. Beyond proliferation, the mitotic advance was associated with higher grade in renal clear cell carcinoma, late stage, whole genome doubling and aneuploidy; Horvath’s clock was not associated with grade or stage and was inversely associated with genome instability. Horvath’s clock read older in renal clear cell and thyroid tumors and younger in four other cancer types, and above chronological age in 98.1% of pediatric tumors. Apart from a HypoClock association reflecting the balance of hyper- and hypomethylation, neither clock family tracked the heterogeneity of methylation change in single region samples, and neither changed consistently at relapse. Female tumors showed X chromosome enrichment of methylation change independent of copy number. Mitotic and chronological clocks therefore record different properties of a tumor: replicative age records methylome remodelling and genome instability, whereas chronological age is specific to cancer type.

## Introduction

Epigenetic clocks, multivariate estimators of age built from DNA methylation (DNAm) at selected CpGs, have become widely used biomarkers of biological ageing^1,2^. Applied to tumors, they are often read as a measure of how far a tumor has aged relative to its host, a view motivated by the shared hallmarks of ageing and cancer^3,4^ and by the long standing hypothesis that a marker of ageing should track the burden of cellular dysregulation in a tumor^5^. Two families of clocks, however, rest on different principles. Chronological clocks were obtained by penalised regression of DNAm against calendar age without a mechanistic prior, from an early predictor of 88 CpGs^6^ to Hannum’s 71 CpG blood clock^7^ and Horvath’s 353 CpG pan tissue clock^8^, and second generation estimators such as PhenoAge and GrimAge were subsequently trained against phenotypic and mortality outcomes^9,10^. Mitotic clocks were instead constructed to count cell divisions, using CpGs that gain methylation at Polycomb targets with each division (epiTOC1 and epiTOC2)^11,12^, loss of methylation at solo WCGW sites in late replicating domains (HypoClock)^13^, or promoter CpGs that are unmethylated in fetal tissue and gain methylation with population doublings but not with the passage of time alone (stemTOC)^14^. Horvath’s clock is therefore not a mitotic clock, and its acceleration measure is not raised in precancerous lesions in which mitotic clocks are^11^.

The rationale for mitotic clocks derives from the finite fidelity with which DNAm is copied at each division. Stochastic epimutations accrue in proportion to replicative history rather than chronological time, so that methylation drift scales inversely with lifespan across mammals^15^ and shared methylation states among cells record their common ancestry, a principle used to trace stem cell lineages in human colon crypts^16^ and extended into fluctuating methylation clocks that resolve stem cell dynamics in normal tissue^17^ and clonal evolution in cancer^18^. Loss of epigenetic fidelity across successive divisions has itself been proposed as a cause of ageing, and experimentally induced loss of fidelity accelerates clock output^19^. Applied to a tumor, this reasoning suggests that the heterogeneity of methylation change between tumor and matched normal tissue might carry information about the replicative history of the malignant population. Methylation derived trees reproduce the topology of mutation derived phylogenies in multiregion and longitudinal tumor samples^20^, and multiregion profiling of lung cancer shows that methylation change cooperates with genomic alteration during tumor evolution^21^. Bulk tumor methylomes, however, are mixtures of malignant, stromal and immune cells, and both mitotic age estimates and the magnitude of tumor methylation change scale with tumor purity^14,22^, so any relation between clock readings and heterogeneity must be assessed against composition.

The behaviour of chronological clocks in tumor tissue has been described across The Cancer Genome Atlas (TCGA). Horvath reported that tumor DNAm age departs markedly from chronological age and from tumor adjacent tissue^8^, Lin and Wagner found that age associated methylation in 25 cancer types hardly reflects patient age but is coherently modified^23^, and comparisons across clock families have found Horvath’s acceleration measure to be among the least consistent markers of tumor state^11,24^. In parallel, mitotic clocks have been found to be accelerated in cancer and in precancerous lesions^11,12,14^. Most of these comparisons reference tumor age against chronological age or against unpaired normal samples, which conflates tumor intrinsic change with inter individual variation in baseline epigenetic age, and few have compared both clock families within the same patients. Childhood tumors offer a complementary setting in which the chronological reference spans only a few years, but they are rarely accompanied by banked normal tissue.

Here we compare four mitotic clocks with Horvath’s clock in 506 matched pairs of primary tumor and normal tissue from 13 adult solid cancers and in 1,731 pediatric tumors spanning 24 entities. We show that mitotic age is advanced in the tumor relative to its own normal tissue in every adult cancer type, independently of purity and composition, whereas Horvath’s clock moves in a direction that depends on cancer type in adults and is near uniformly advanced in children. By partitioning the mitotic advance between proliferation, composition, global methylation change and genome instability, we find that current proliferation accounts for only a small share of it, although the advance carries information on tumor grade, stage and aneuploidy that proliferation does not. By relating clock differences to the heterogeneity of methylation change at 500 bp resolution in each patient, and following pediatric tumors from primary to relapse, we find that neither clock family tracks clonal diversification in bulk tissue. Finally, we show that the apparent enrichment of methylation change on the X chromosome is confined to female tumors and persists in the absence of X copy number change.

## Results

### A matched design across adult and pediatric tumors

To compare clocks within patients, we rebuilt the adult cohort from TCGA HumanMethylation450 (HM450) beta value files so that each patient contributed exactly one primary tumor and one solid tissue normal sample (“Methods” section). Of 545 patients with tumor and normal files, 531 satisfied this rule, and the 506 pairs from the 13 cancer types with at least ten pairs formed the main analysis (Fig. 1a,b, Table 1, Supplementary Table 1). Pediatric tumors were drawn from the CGCI, TARGET, HCMI and TCGA projects; 1,731 tumors from 1,721 patients had a recorded age at diagnosis (median 5.4 years), of which 1,685 belonged to 24 entities with at least ten tumors. No matched normal tissue was available in the pediatric cohort, so pediatric clocks were referenced to chronological age. Pediatric samples had been profiled on HM450, EPIC and EPICv2 arrays, and platform was largely determined by source project and hence by entity; all pediatric comparisons of mitotic clocks were therefore made within platform, using CpGs shared by all three arrays (“Methods” section).

**Fig. 1.**
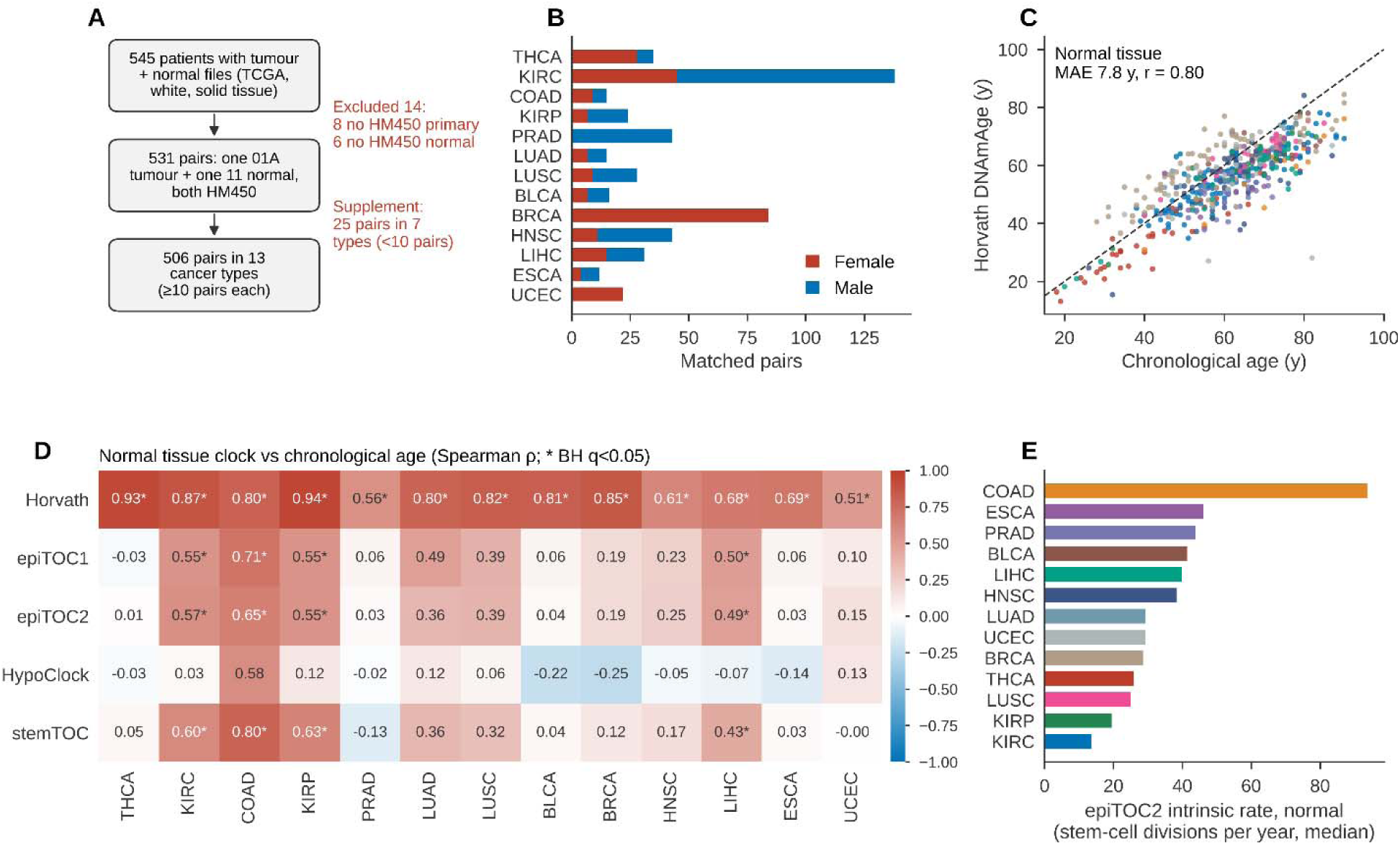
| Cohort and clock calibration in normal tissue. **a**, Cohort flow from 545 patients with tumor and normal files to 531 pairs, and to 506 pairs in 13 cancer types with at least ten pairs. **b**, Matched pairs per cancer type by donor sex. **c**, Horvath DNAmAge against chronological age in adult normal tissue; mean absolute error 7.8 years, r = 0.80. **d**, Spearman correlation of each clock with chronological age in normal tissue, per cancer type. **e**, epiTOC2 intrinsic division rate per year in normal tissue.

**Table 1.**
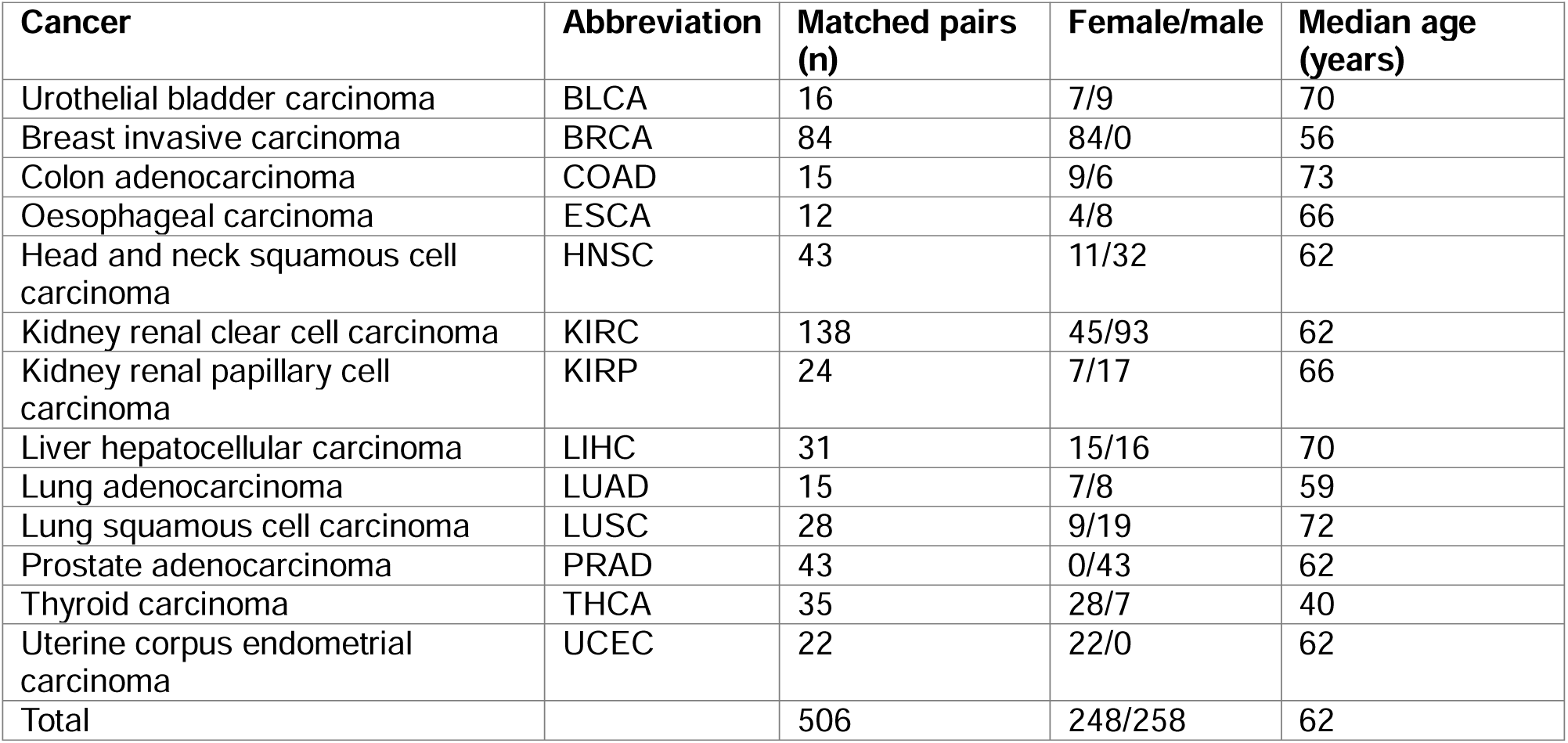
| Adult cancer types in the main analysis. Number of patients contributing one primary tumor and one matched normal sample, donor sex and median age at diagnosis. Seven cancer types with fewer than ten pairs (25 pairs) are listed in Supplementary Table 2.

### Mitotic age in normal tissue reflects tissue turnover more than calendar age

We reasoned that in normal tissue a chronological clock should track calendar age closely, whereas a mitotic clock should track it only to the extent that stem cell turnover is constant over life, and should differ between tissues according to their division rates. Consistent with this, Horvath’s clock reproduced chronological age in adult normal tissue with a mean absolute error of 7.8 years (r = 0.80; Fig. 1c), whereas the four mitotic clocks were more weakly coupled to age. Pooled within cancer type, the correlation of epiTOC1, epiTOC2 and stemTOC with age was 0.23 to 0.26, and that of HypoClock was absent (ρ = −0.03); correlations reached significance within individual cancer types only in COAD, KIRC, KIRP and LIHC (ρ 0.4 to 0.8, q < 0.05; Fig. 1d, Supplementary Table 3), the four cancer types in which an independent mitotic age estimator also tracks age in adjacent normal tissue^25^. The intrinsic division rate estimated by epiTOC2^12^ differed by almost sevenfold between normal tissues, from 13.6 divisions per year in kidney (KIRC) and 19.5 in KIRP to 44 in prostate, 46 in oesophagus and 93.8 in colon (Fig. 1e), in line with the high turnover of colonic epithelium^12,14^, and the ranking of cancer types by intrinsic rate was preserved in tumors (ρ = 0.76).

The same pattern was evident in childhood tumors, in which the chronological range is narrow. Within entity and platform, epiTOC1, epiTOC2 and stemTOC rose with the age of the child in 32 of 112 tests (q < 0.05; ρ 0.2 to 0.85), for example in malignant rhabdoid tumor (ρ = 0.73), neuroblastoma, Wilms tumor, ependymoma and medulloblastoma, whereas HypoClock showed no association with age in any entity (Supplementary Table 4b), mirroring its behaviour in adult normal tissue. Horvath DNAmAge in pediatric tumors likewise retained an appreciable association with chronological age (Pearson r = 0.54, Spearman ρ = 0.60) while sitting systematically above it, as described below. Thus, in both cohorts the hypermethylation based mitotic clocks carry an age signal that is weaker and more tissue dependent than that of the chronological clock, as expected of division counters.

### Mitotic age is advanced in tumors of every adult cancer type

We next compared each tumor with its own normal tissue. All four mitotic clocks read higher in the tumor in all 13 cancer types (paired Wilcoxon test, q < 0.01 for every clock and cancer type), and no cancer type showed a negative median difference (Fig. 2a,b, Supplementary Table 5). For stemTOC, 97% of tumors exceeded their matched normal sample, with q < 0.001 in 12 cancer types and q < 0.01 in ESCA. THCA showed the weakest effect, with 71 to 83% of pairs higher in the tumor depending on the clock. The four clocks largely carried a shared signal: tumor minus normal differences in epiTOC1 and epiTOC2 were nearly identical (ρ = 0.99), stemTOC correlated with both at ρ of about 0.8, and HypoClock correlated less strongly (ρ 0.4 to 0.5; Fig. 2d). This near universal advance replicates the acceleration of mitotic clocks reported in comparisons with tumor adjacent tissue^11,12,14,25^ and establishes it within individual patients, providing a positive control for the matched design.

**Fig. 2.**
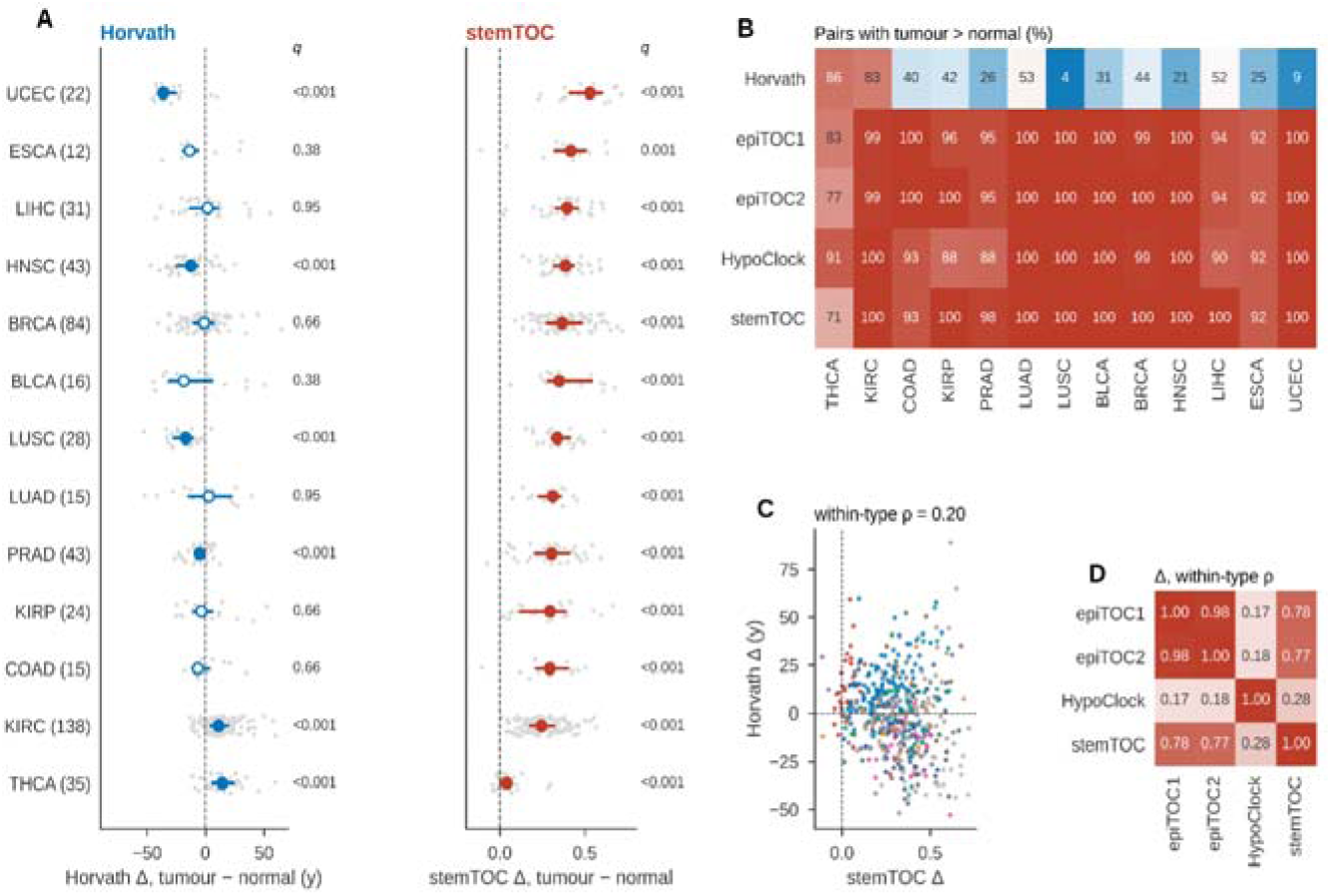
| Tumor minus normal differences in mitotic and chronological clocks. **a**, Median and IQR of tumor minus normal differences in Horvath DNAmAge and stemTOC per cancer type, ordered by median stemTOC difference; filled markers, q < 0.05; open markers, not significant (paired Wilcoxon test with Benjamini and Hochberg correction). **b**, Percentage of pairs in which the tumor exceeds its matched normal sample, for five clocks and 13 cancer types. **c**, Per patient Horvath difference against stemTOC difference; ρ = 0.20 within cancer type. **d**, Within cancer type correlation between tumor minus normal differences of the four mitotic clocks.

### Chronological clock readings of tumors are bidirectional in adults and unidirectional in children

Horvath’s clock behaved differently. In adult tumors its mean absolute error against chronological age rose to 17.6 years and its correlation with age fell to 0.28, in agreement with the observation that age associated methylation hardly reflects patient age in cancer^23^. Relative to matched normal tissue, tumor DNAmAge was higher in KIRC (median +11.3 years) and THCA (+14.5 years), and lower in UCEC (−36.0 years), LUSC (−16.9 years), HNSC (−12.5 years) and PRAD (−5.0 years; all q < 0.001), while the remaining seven cancer types did not differ significantly from zero despite wide spreads in BLCA (median −18.4 years) and ESCA (−13.5 years) (Fig. 2a, Supplementary Table 6a). Across the cohort only half of tumors read older than their matched normal sample, compared with 97% for stemTOC (Fig. 2b). This dependence on cancer type has been described previously^8,23^, and the deceleration in UCEC agrees with the deceleration relative to chronological age reported for the TCGA endometrial cohort^26^. Per patient differences in Horvath DNAmAge and stemTOC were only weakly correlated (ρ = 0.20 within cancer type; Fig. 2c), indicating that the two clock families report largely different quantities in the same pair of samples.

In childhood tumors, where acceleration could be referenced only to chronological age, the direction of change was near uniform. A total of 1,698 of 1,731 tumors (98.1%) returned a DNAmAge above the patient’s chronological age, with a median discrepancy of +13.7 years (IQR 6.6 to 26.3; Wilcoxon signed rank test, P = 6.1 × 10^−280^), so that the typical childhood tumor, at a median age of 5.4 years, returned a clock reading of 20.3 years. The magnitude of this advancement differed markedly between entities (Fig. 3a,b; Kruskal Wallis test, P = 2.4 × 10^−74^), with medians spanning approximately sevenfold, from +5.6 years in oligoastrocytoma and mixed glioma, +6.0 in clear cell sarcoma of the kidney and +6.2 in retinoblastoma, to +23.8 in osteosarcoma, +25.9 in ependymoma, +33.5 in rhabdomyosarcoma and +37.2 in synovial sarcoma. No entity returned a negative median, and the paired age distributions show that the displacement is a shift of the whole distribution rather than the influence of a tail (Fig. 3a). Ordered by anatomical site the same gradient was apparent (Fig. 3c; P = 4.5 × 10^−32^), with ocular (+34.3 years), extra limb bone (+30.1), soft tissue (+24.8) and limb bone (+20.2) tumors at one extreme, brain tumors intermediate (+15.3), and retroperitoneal (+10.2), renal (+7.7) and adrenal (+7.2) tumors at the other; within site, DNAmAge remained positively correlated with chronological age in seven of eight sites.

**Fig. 3.**
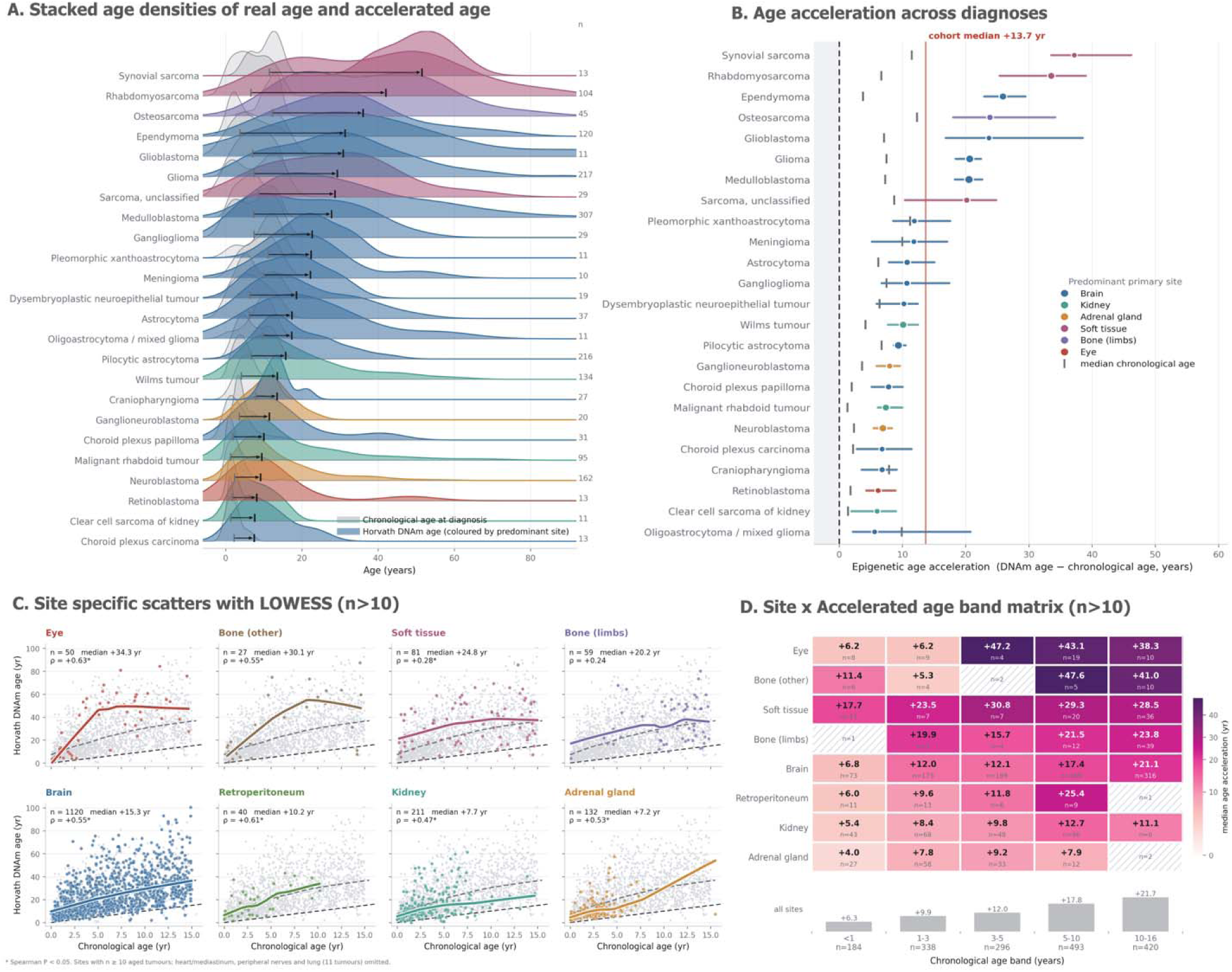
| Horvath DNAmAge in pediatric tumors. **a**, DNAmAge and chronological age at diagnosis by entity. **b**, Acceleration (DNAmAge minus chronological age) by entity. **c**, Acceleration by anatomical site. **d**, Acceleration by age band. Entities and sites with at least ten tumors are shown.

Acceleration also increased with the age of the child, from a median of +6.3 years in infants under one year to +21.7 years at 10 to 16 years (Fig. 3d). This gradient is at least partly structural: because DNAmAge cannot fall below zero, the attainable negative acceleration is bounded by the patient’s own age, and the metric correlates positively with chronological age across the cohort (ρ = 0.35). The site ranking was nonetheless reproduced within age bands from three years upward, where ocular, bone and soft tissue tumors exceeded renal and adrenal tumors in every band; below three years, cells of four to nine tumors did not reproduce the overall ranking. Because platform and entity are largely confounded in this cohort, we asked whether the entity differences held for mitotic clocks compared within platform. Platform effects were material, with mitotic age in neuroblastoma differing between HM450, EPIC and EPICv2 samples (epiTOC2 P = 0.001, stemTOC P = 0.006) and a fivefold difference in epiTOC2 between Wilms tumors on HM450 and nephroblastomas on EPIC (Supplementary Fig. 1). Within platform, mitotic age differed strongly between entities (Kruskal Wallis P < 10^−10^ for all clocks): among 23 entities profiled on EPIC it was highest in embryonal and alveolar rhabdomyosarcoma, rhabdoid tumors and sarcoma and lowest in dysembryoplastic neuroepithelial tumor, pilocytic astrocytoma and astrocytoma, and among five entities on HM450 it was lowest in Wilms tumor and highest in osteosarcoma and clear cell sarcoma of the kidney (Supplementary Table 4a). Mesenchymal tumors thus ranked high on both clock families, whereas clear cell sarcoma of the kidney ranked among the least advanced entities by Horvath’s clock and among the most advanced by mitotic clocks, again showing that the two families can order the same tumors differently.

### Clock differences are robust to tumor purity and cell type composition

Because bulk tumor and normal tissue differ in composition, a tumor minus normal clock difference could in principle arise from cell mixture alone. To assess this, we estimated epithelial, fibroblast and immune fractions in every adult sample with EpiDISH^27^ and obtained tumor purity from the TCGA consensus estimate (CPE)^22^. EpiDISH epithelial fractions in tumors correlated with CPE in 10 of 12 cancer types (ρ 0.52 to 0.83; Fig. 4b, Supplementary Table 7); the exceptions were LIHC, in which deconvolution failed in the absence of a hepatocyte reference, and PRAD, in which all tumors had a CPE of at least 0.75. Tumors of UCEC, THCA, LUSC and BLCA were enriched for epithelial cells relative to their normal counterparts (median increase 0.13 to 0.29), and the adipocyte fraction in BRCA fell from 0.61 in normal tissue to 0.33 in tumors (Fig. 4a, Supplementary Table 8). Restricting the analysis to tumors with CPE of at least 0.6 left every mitotic clock significant in every cancer type tested (Fig. 4c). Adjusting for changes in fibroblast and immune fractions left the same six cancer types significant for Horvath’s clock, in the same direction (for example, KIRC +12.8 years and UCEC −27.8 years; Supplementary Table 9). Under this adjustment the mitotic clock difference in THCA was attenuated; however, the composition coefficients were mostly not significant and most THCA pairs differed in epithelial fraction by more than 0.1, so the adjusted estimate extrapolates beyond the observed data. We therefore regard the THCA mitotic signal as robust to purity restriction and attenuated only under extrapolated adjustment.

**Fig. 4.**
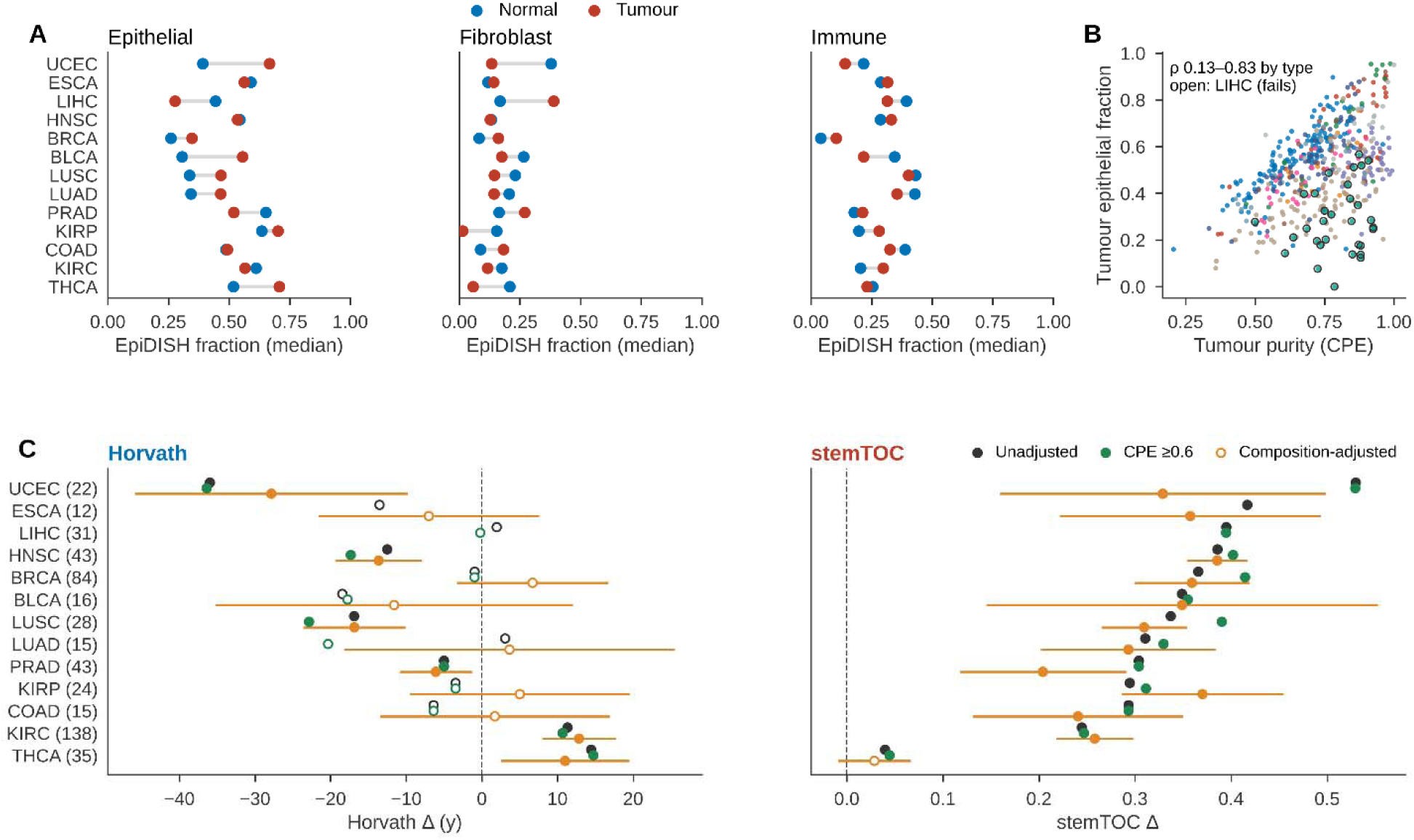
| Tumor purity and cell type composition. **a**, EpiDISH epithelial, fibroblast and immune fraction in normal and tumor tissue per cancer type. **b**, Tumor epithelial fraction against CPE per cancer type; deconvolution fails in LIHC. **c**, Tumor minus normal differences in Horvath DNAmAge and stemTOC, unadjusted, restricted to CPE of at least 0.6, and adjusted for composition.

### Proliferation accounts for a small part of the mitotic advance

A tumor minus normal mitotic clock difference may integrate current proliferation, the replicative history of the cell of origin and global remodelling of the methylome. Earlier work related tumor mitotic clock readings separately to an expression based mitotic index^11^ and to the fraction of the putative cell of origin^14^, in tumor samples only. To separate these components within patients, we scored proliferation in 503 tumors and 273 matched normal samples with RNA sequencing data, using the nine gene mitotic index of Yang et al.^11^ as the primary score and a Hallmark G2M checkpoint score^28^ for sensitivity analyses (median within type ρ between scores 0.85; Supplementary Table 10, Supplementary Table 11). Proliferation was higher in the tumor than in matched normal tissue in at least 87% of pairs in 12 of 13 cancer types and lowest in THCA (78%), paralleling the weakest mitotic advance. In tumor tissue alone, stemTOC correlated positively with proliferation in 12 of 13 cancer types (ρ 0.07 to 0.59), reproducing the published direction^11^, whereas Horvath DNAmAge correlated negatively in 12 (Supplementary Table 12b).

In a pooled model of the tumor minus normal clock difference on tumor proliferation, adjusted for purity, changes in epithelial, fibroblast and immune fractions, the signed global Δβ, age, sex and cancer type (480 pairs), proliferation was positively associated with all four mitotic clocks (standardised β 0.16 for epiTOC1, epiTOC2 and HypoClock, q ≤ 0.011; 0.26 for stemTOC, q = 9 × 10^−6^) and not with Horvath’s clock (β = −0.04, q = 0.51; Fig. 5a, Supplementary Table 13). These associations persisted with the G2M score (β 0.22 to 0.30), with the tumor minus normal proliferation difference in 257 complete pairs (β 0.19 to 0.28) and, for stemTOC and HypoClock, in tumors with a CPE of at least 0.6. They rested largely on KIRC, however. Within cancer type, proliferation was significant only in KIRC (β 0.39 to 0.45 for epiTOC1, epiTOC2 and stemTOC, q = 0.037; Fig. 5c, Supplementary Table 12a), and after exclusion of KIRC the association remained for stemTOC (β = 0.19, q = 0.003) but not for epiTOC1 or epiTOC2 (β = 0.05, q = 0.51). Partitioning the variance of the clock difference in 465 pairs showed that proliferation accounted for 2.4 to 3.0% for epiTOC1, epiTOC2 and HypoClock and 6.7% for stemTOC, whereas the global Δβ accounted for 10.9 to 25.2% and composition for 18.0% of the stemTOC and 10.2% of the HypoClock difference; whole genome doubling and aneuploidy together contributed at most 2.7%, and no block explained more than 1.3% of the Horvath difference (Fig. 5b, Supplementary Table 14). Telomere length ratios between tumor and normal tissue^29^, available for 361 pairs and shorter in the tumor in 55 to 100% of pairs per cancer type, explained at most 2% of any mitotic clock difference except HypoClock (6.3%; Supplementary Table 15).

**Fig. 5.**
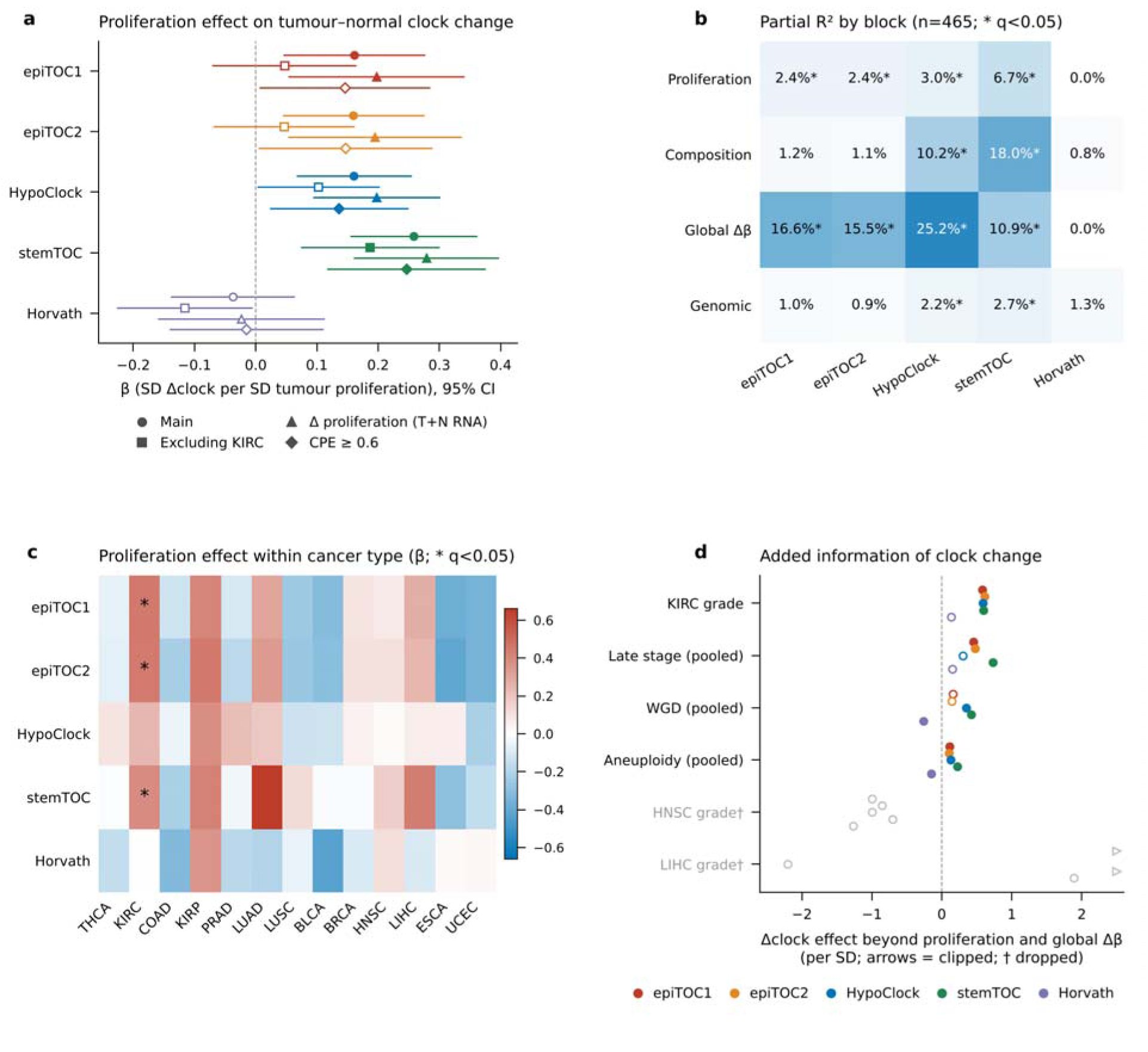
| Proliferation, composition and genome instability in tumor minus normal clock differences. **a**, Standardised association of tumor proliferation (nine gene mitotic index) with the tumor minus normal difference of each clock, adjusted for CPE, changes in epithelial, fibroblast and immune fractions, signed global Δβ, age, sex and cancer type, in the main model, excluding KIRC, using the tumor minus normal proliferation difference, and restricted to CPE of at least 0.6; bars, 95% CI; filled markers, q < 0.05. **b**, Drop one partial R² of the clock difference for proliferation, composition, global Δβ and genomic (aneuploidy and whole genome doubling) blocks in 465 pairs; *, q < 0.05. **c**, Proliferation coefficient within cancer type, ordered by median stemTOC difference; *, q < 0.05. **d**, Association of the clock difference with KIRC grade, late stage, whole genome doubling and aneuploidy beyond proliferation, composition and global Δβ; filled markers, q < 0.05; grey, HNSC and LIHC grade, not interpreted owing to separation.

The mitotic advance nonetheless carried information that proliferation did not. When the clock difference was added to models of clinical and genomic features that already contained tumor proliferation, composition, global Δβ, age and sex (Supplementary Table 16), all four mitotic clocks were associated with higher grade in KIRC (β 0.58 to 0.61 per standard deviation, q = 0.004; n = 134), and epiTOC1, epiTOC2 and stemTOC with late stage across KIRC, BRCA, KIRP and THCA (β 0.45 to 0.73, q ≤ 0.005; n = 269), whereas Horvath’s clock was associated with neither (q ≥ 0.27; Fig. 5d, Supplementary Table 17). stemTOC and HypoClock were associated with whole genome doubling (β 0.42 and 0.35, q ≤ 0.010; n = 466) and all four mitotic clocks with aneuploidy (β 0.11 to 0.22, q ≤ 0.032; n = 395), while Horvath’s clock was associated with both in the opposite direction (β = −0.26, q = 0.033; β = −0.15, q = 0.006), in keeping with the low Horvath age acceleration of tumors with many somatic mutations^8^. The tumor minus normal mitotic advance therefore records global methylation change and tissue composition first, contains a proliferative component that is general only for stemTOC, and tracks grade, stage and genome instability in the direction opposite to that of the chronological clock.

### Clock readings do not track the heterogeneity of methylation change

Because the mitotic advance was dominated by the global magnitude of methylation change, we asked whether, beyond that magnitude, tumors with a greater advance show more heterogeneous methylation change, as would be expected if heterogeneity records replicative history. To this end, we summarised tumor minus normal differences in mean beta value (Δβ) over probe clusters of consecutive CpGs separated by no more than 500 bp, a scale matched to the decay of methylation correlation^30,31^, yielding about 54,000 units per pair (“Methods” section, Supplementary Table 18). Mixture modelling of |Δβ| identified a single component in 513 of 531 pairs (97%), so discrete tiers of methylation change, as would be expected from clonal and subclonal events, could not be resolved from single region bulk samples (Fig. 6a, Supplementary Table 19). We therefore summarised heterogeneity by the discordance of methylation change, the share of changed units whose direction opposes the patient’s majority direction (Fig. 6b, Supplementary Table 20), the only candidate metric not dominated by the overall magnitude of Δβ (Supplementary Table 21). Discordance was independent of tumor purity and of changes in fibroblast and immune fractions (all q ≥ 0.34), whereas magnitude summaries such as the standard deviation of Δβ and the fraction of changed units scaled with purity, as expected from dilution by non tumor cells, and the signed mean Δβ fell with stromal gain (Fig. 6c, Supplementary Table 22).

**Fig. 6.**
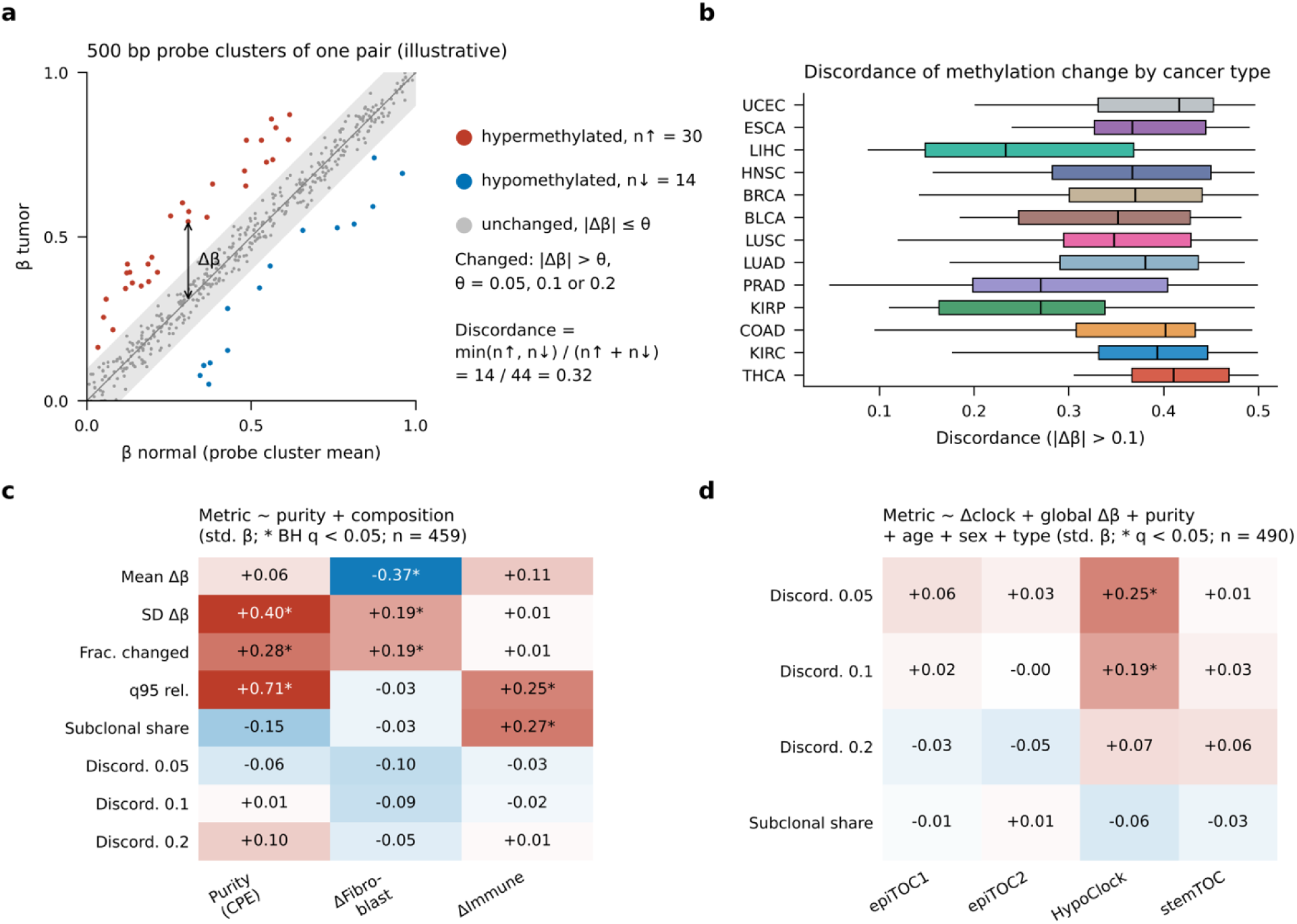
| Heterogeneity of methylation change. **a**, Schematic of the 500 bp probe cluster analysis. **b**, Discordance of methylation change per cancer type. **c**, Pooled association of heterogeneity and magnitude metrics with CPE and with changes in fibroblast and immune fractions. **d**, Association of tumor minus normal clock differences with heterogeneity metrics after adjustment for the signed and absolute mean Δβ, spread of Δβ, CPE, age, sex and cancer type.

In a model accounting for the signed and absolute mean Δβ, the spread of Δβ, purity, age, sex and cancer type, no clock was associated with any measure of heterogeneity except for a positive association between HypoClock and discordance (β = 0.020, q = 0.0065; Fig. 6d, Supplementary Table 23). HypoClock is built from hypomethylation at solo WCGW sites^13^ and tracked the signed mean Δβ (ρ = −0.51), so this association reflects the balance between hypermethylation and hypomethylation rather than heterogeneity as such. Associations of epiTOC1, epiTOC2 and stemTOC with discordance disappeared once the signed mean Δβ was included. At 5 Mb resolution, as in the original window based analysis, discordance was positively associated with epiTOC and stemTOC, but this did not replicate at 500 bp and we regard it as an effect of resolution (Supplementary Fig. 2, Supplementary Table 24). The 5 Mb summary did, however, reproduce the dominant direction of change in each cancer type: KIRP and PRAD were dominated by hypermethylation (84 to 90% of events), whereas KIRC, THCA, BLCA and LUSC were dominated by hypomethylation, so that the two renal cancers arising from the same organ diverged in the dominant direction of methylation change (Supplementary Table 25).

A cross sectional pair cannot show whether clock readings advance as a tumor evolves. The pediatric cohort included ten patients with paired primary and relapse tumors, five Wilms tumors, four neuroblastomas and one ganglioneuroblastoma, all profiled on HM450, which provide a limited longitudinal test (Fig. 7). Seven of the ten relapse tumors returned a higher Horvath DNAmAge than the corresponding primary, with a median within patient change of +1.8 years (IQR −4.2 to +8.3), but the shift did not differ from zero (Wilcoxon signed rank test, P = 0.38), and the change spanned a 27 year range, with the clock running backwards at relapse by 6.1, 6.6 and 8.5 years in two Wilms tumors and one neuroblastoma. Mitotic clocks showed no consistent change either, with three to five of ten relapse tumors higher than the primary (exact P = 0.16 to 0.92 across clocks), unchanged when one relapse sampled from bone marrow was excluded (Supplementary Table 4c,d). With ten pairs these tests are underpowered, but together with the adult analysis they provide no evidence that either clock family tracks the course of tumor evolution in bulk samples.

**Fig. 7.**
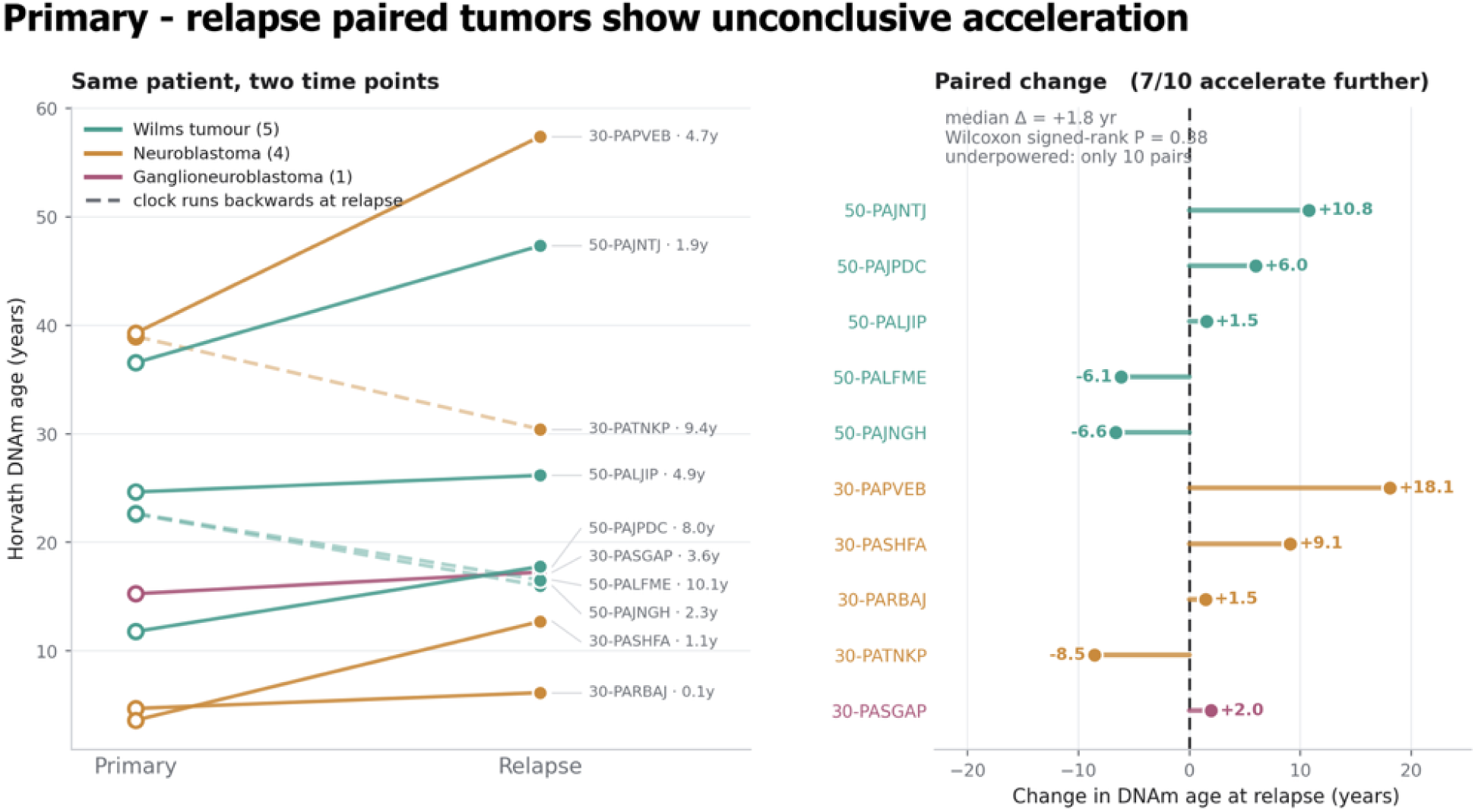
| Clock readings in paired primary and relapse pediatric tumors. Horvath DNAmAge in primary and relapse tumors from ten patients (five Wilms tumors, four neuroblastomas and one ganglioneuroblastoma).

### X chromosome enrichment of methylation change is female specific

In the original window based analysis, the share of methylation events on the X chromosome exceeded the mean autosomal share in 19 of 20 cancer types. Because this comparison is partly driven by chromosome size, we normalised the rate of changed units on the X by the autosomal rate in each patient. After normalisation, X enrichment was significant in four of 13 cancer types, strongest in THCA (ratio 1.82) and BRCA (1.58, q = 1 × 10^−7^) and modest in KIRC (1.13) and PRAD (1.09, q = 0.025) (Fig. 8a, Supplementary Table 26, Supplementary Table 27). Stratifying by sex showed that enrichment was confined to female donors, reaching 2.05 in THCA (q = 3 × 10^−8^), 1.58 in BRCA, 1.34 in KIRC (q = 0.001) and 1.29 in LIHC (q = 0.04), whereas in men it was close to one in KIRC (1.01), HNSC (1.04) and LUSC (1.00; Supplementary Table 28). In female tumors the direction of enrichment differed between cancer types (Fig. 8b): it was driven by hypomethylation in BRCA (2.76), THCA (2.11), HNSC (2.62) and LIHC (1.53), the pattern expected from erosion of methylation on the inactive X, and by hypermethylation in KIRC (2.18) and UCEC (2.11).

**Fig. 8.**
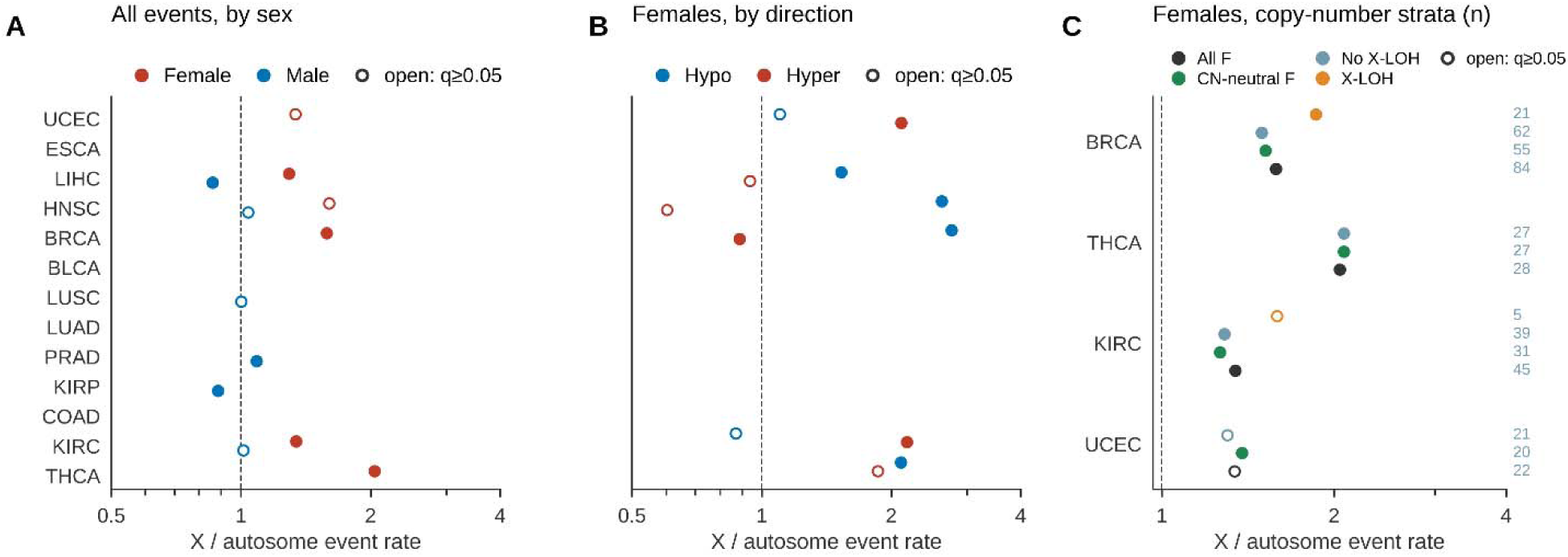
| X chromosome enrichment of methylation change. **a**, Ratio of the rate of changed units on the X chromosome to the autosomal rate, by cancer type and donor sex. **b**, Female enrichment separated into hypomethylation and hypermethylation. **c**, Female enrichment in BRCA, THCA, KIRC and UCEC b X copy number state.

Loss or gain of an X copy would itself produce a large Δβ across the chromosome. Female enrichment, however, persisted in tumors without X copy number change, at 1.52 in BRCA (hypomethylation 2.56), 2.08 in THCA, 1.26 in KIRC (hypermethylation 2.29) and 1.38 in UCEC (hypermethylation 2.68, q = 0.006) (Fig. 8c, Supplementary Table 29). In BRCA, tumors with X loss of heterozygosity showed stronger enrichment (1.86, n = 21) than those without (1.50, n = 62), indicating that copy loss contributes to the signal but does not account for it. In male tumors we observed an excess of hypermethylation on the X in KIRC (1.49, q = 6 × 10^−10^) and HNSC (1.41), which remains unexplained; male X copy number estimates were not interpretable and could not be used to resolve it (Supplementary Table 30).

## Discussion

We have here compared mitotic and chronological epigenetic clocks within patients, by referencing each adult tumor to its own normal tissue, and have extended the comparison to childhood tumors and to paired primary and relapse samples. Three main insights emerge. First, replicative age, as measured by four mitotic clocks, is advanced in tumors of every adult cancer type examined, independently of purity and composition, and this advance reflects global methylation remodelling and composition more than current proliferation. Second, Horvath’s chronological clock reads tumors in a direction that depends on cancer type in adults and is near uniformly advanced in children, and its per patient readings are only weakly related to mitotic age. Third, neither clock family tracks the heterogeneity of methylation change in bulk tissue, either across patients or from primary to relapse.

The near universal advance of mitotic clocks agrees with earlier evidence that epiTOC, epiTOC2, HypoClock and stemTOC are accelerated in cancer and precancerous lesions^11,12,14^, and the matched design shows that this advance is present within each patient against that patient’s own normal tissue. It is worth stressing that mitotic age estimates scale with the tumor cell of origin fraction^14^, so an advance in bulk tumor tissue could partly reflect enrichment of epithelial cells. Restriction to tumors with a purity of at least 0.6 left the advance significant in every cancer type tested, however, which argues against dilution by non tumor cells as its main explanation. The intrinsic division rates estimated in normal tissue, highest in colon and lowest in kidney, are in line with the tissue ranking reported for epiTOC2^12^, and the preservation of this ranking in tumors suggests that tumor mitotic age inherits part of its value from the tissue of origin. THCA showed the weakest advance and the smallest excess of tumor over normal proliferation, consistent with the slow turnover of thyroid follicular epithelium. In children, the rise of mitotic age with age within entities, and the absence of such a rise for HypoClock, reproduce in tumor tissue the behaviour of these clocks in adult normal tissue and are consistent with the view that clocks anchored on methylation gains are better proxies of mitotic age than those based on hypomethylation^12,14^.

The joint model qualifies what this advance measures. Tumor mitotic clock readings have been related to an expression based mitotic index^11^ and to the fraction of the presumed cell of origin^14^ in separate analyses of tumor samples. Estimated within patients and in the same model, these components were unequal: global methylation change and composition accounted for most of the explained variance and current proliferation for 2 to 7%, with a cross cancer association that held for stemTOC but, for epiTOC1 and epiTOC2, rested on clear cell renal carcinoma. The share attributed to global Δβ is partly mechanical, because clocks and global Δβ summarise the same tumor and normal methylomes, but its size shows that a mitotic clock difference in bulk tissue is dominated by the remodelling that accompanies transformation rather than by the division rate at sampling. The associations of HypoClock with proliferation, genome doubling and aneuploidy agree with the dependence of solo WCGW hypomethylation on cell division and its link with copy number load^13,32,33^, and the association of stemTOC with composition agrees with its dependence on the cell of origin fraction^14^. That the mitotic advance nevertheless tracked grade, stage and aneuploidy beyond proliferation, while Horvath’s clock tracked genome instability inversely, in line with its lower acceleration in tumors with high somatic mutation burden^8^, shows that the two clock families carry distinct and partly opposed information on tumor state.

The behaviour of Horvath’s clock in adult tumors reproduces earlier reports. Horvath described marked variation in tumor DNAm age across cancer types^8^, Lin and Wagner showed that age associated methylation in cancer is coherently modified rather than tied to patient age^23^, and the deceleration we observe in UCEC agrees with the deceleration relative to chronological age reported in the TCGA endometrial cohort, in which high deceleration marked an aggressive, immune excluded phenotype^26^. That this clock tracked chronological age well in normal tissue, including breast and endometrium in which pan tissue clocks are known to be less accurate^8,34^, but lost most of its relation to age in tumors, and that its readings were only weakly related to mitotic age in the same pairs, indicates that the two families capture different processes: a chronological clock trained on age associated CpGs, whose reading in tumors depends on how those CpGs are remodelled in each tissue, and a division counter that rises with replicative history in every tissue examined. The pediatric results fit this picture. The mean absolute discrepancy between Horvath DNAmAge and chronological age in pediatric tumors, 18.5 years, exceeds the clock’s error in adult normal tissue and matches its error in adult tumors (17.6 years), so it is consistent with the loss of age correspondence that the clock shows in malignant tissue generally, and we do not interpret the pediatric medians as measurements of biological age in years. What distinguishes the pediatric observation is its consistency of direction. Part of this unidirectionality is a property of the scale: in an adult, a tumor reading 17 or 36 years below its matched normal sample, as observed in LUSC and UCEC, is readily expressible, whereas in a child with a median age of 5.4 years the same deceleration cannot be expressed, because DNAmAge is bounded below by zero, and the most decelerated pediatric tumor sits 8.8 years below chronological age. The ranking of pediatric entities, with mesenchymal tumors high on both clock families but clear cell sarcoma of the kidney discordant between them, further shows that the two families can order the same tumors differently.

The absence of an association between clock readings and the heterogeneity of methylation change deserves careful interpretation. We had reasoned that if heterogeneity of methylation change records replicative history, tumors with a larger mitotic advance should show more heterogeneous change. After accounting for the overall magnitude and direction of change, purity, age and sex, no mitotic clock was associated with heterogeneity, and the single association that remained, between HypoClock and discordance, is most simply explained by HypoClock’s dependence on hypomethylation in late replicating domains^13^, which ties it to the balance of gains and losses rather than to their dispersion. Two features of the data limit what this null result can show. First, a single bulk sample from one region per tumor did not resolve discrete tiers of methylation change in 97% of pairs, so clonal and subclonal events cannot be separated by the variance structure of Δβ alone. Second, an association seen at 5 Mb did not replicate at 500 bp, which shows that coarse windows can produce structure that is not present at the scale of comethylation. The relapse pairs, although few, point in the same direction. Multiregion and longitudinal methylation data, in which clonal changes are identified by their presence across regions or time points^18,20,21^, are therefore the appropriate setting in which to test whether replicative history and the clonal architecture of methylation change are related. The divergence in dominant direction of change between KIRC and KIRP, two cancers of the same organ, nonetheless shows that the direction of methylation change in bulk tumors is a cancer type specific feature that is reproducible across resolutions.

The X chromosome finding changed in character once the analysis was normalised and stratified. The apparent enrichment in 19 of 20 cancer types was in part a consequence of chromosome size, and after normalisation it was significant in four cancer types and confined to women. In BRCA, THCA, HNSC and LIHC it was driven by hypomethylation and persisted in X copy number neutral tumors, consistent with epigenetic erosion of the inactive X, a recurrent feature of breast cancer^35,36^. A pan cancer measure of defective X inactivation built from TCGA methylation data likewise ranked thyroid and breast cancer among the cancer types with the highest frequency of X reactivation, and found that defective X inactivation, rather than demethylation of genes that escape inactivation, accounted for its association with cancer risk^37^. The hypermethylation driven enrichment in female KIRC and UCEC does not fit this pattern, indicating that more than one process can concentrate methylation change on the X.

It is also important to discuss the limitations of this study. The adult cohort was restricted to white donors, which reduces confounding by ancestry but limits generalisability. Normal samples were tumor adjacent tissue, which may carry field effects that reduce tumor and normal contrasts^24^. Purity estimates were unavailable for ESCA, composition could not be estimated reliably in liver, and composition adjusted estimates rest on extrapolation where tumor and normal composition differ widely. The X chromosome analysis was performed at 5 Mb resolution with about 30 X windows per patient, and copy number was evaluated at the patient level only, because male X copy number estimates were unusable. Proliferation was measured from a single bulk RNA profile per sample, the cross cancer proliferation association of epiTOC1 and epiTOC2 and the clinical associations rest substantially on KIRC, grade could be modelled only in KIRC, and telomere length was estimated mostly from exome sequencing. In the pediatric cohort, platform and entity are largely confounded, no matched normal tissue was available and only ten relapse pairs could be studied.

In summary, our data indicate that tumor epigenetic age measured as replicative history is near uniformly advanced relative to matched normal tissue across adult solid cancers, reflects global methylation remodelling and tissue composition more than current proliferation, and tracks tumor grade, stage and aneuploidy, whereas the chronological clock reading of a tumor depends on its cancer type in adults and is consistently advanced in children. Neither readout tracks the heterogeneity of methylation change in single region bulk samples, so whether replicative age and the clonal architecture of methylation change are related remains to be tested in multiregion data. The female specific, copy number independent enrichment of methylation change on the X chromosome points to erosion of X inactivation in a subset of female tumors and merits sex stratified study of its clinical relevance.

## Methods

### Adult cohort

DNAm beta values were obtained from TCGA through the Genomic Data Commons (GDC) portal. Samples were restricted to white donors with open access solid tissue data profiled on the Illumina HM450 BeadChip. Sample type was read from the TCGA aliquot barcode, and each aliquot was mapped to its GDC beta value file through the GDC application programming interface, yielding 1,062 files with no unmapped and no duplicated aliquots. Each patient contributed one primary tumor (sample type 01, vial A, lowest portion) and one solid tissue normal sample (sample type 11), both profiled on HM450. Metastatic and recurrent tumors (sample types 06, 02 and 05) were excluded, and technical replicates were collapsed to a single sample. Of 545 patients with tumor and normal files, 531 satisfied these criteria; eight lacked an HM450 primary tumor and six an HM450 normal sample (Supplementary Fig. 3, Supplementary Table 1). Donor sex, age and race were retrieved from the GDC clinical records, and sex restricted cancer types were confirmed as female only for BRCA and UCEC and male only for PRAD. Cancer types with at least ten matched pairs entered the main analyses (506 pairs, 13 cancer types; Table 1); the remaining seven types (25 pairs) are reported in Supplementary Table 2 only.

### Pediatric cohort

Pediatric beta matrices were obtained from the GDC portal, drawing on the Cancer Genome Characterization Initiative pediatric brain and solid tumor projects (CGCI/PB), the Therapeutically Applicable Research to Generate Effective Treatments programme (TARGET), the Human Cancer Models Initiative (HCMI) and TCGA. This yielded 2,125 tumor samples from 2,115 patients across 11 anatomical sites, of which 1,714 derived from CGCI/PB, 390 from TARGET and 21 from HCMI, TCGA and one further submission. Samples had been profiled on HM450 (335 samples), EPIC (1,654) and EPICv2 (148); Wilms tumor, neuroblastoma, osteosarcoma, ganglioneuroblastoma and clear cell sarcoma of the kidney were profiled mainly on HM450, and all central nervous system tumors on EPIC. No matched normal tissue was available. Of the 2,125 samples, 394 lacked a recorded age at diagnosis and were excluded from all age referenced analyses, leaving 1,731 tumors from 1,721 patients (median age 5.4 years, IQR 2.6 to 9.9, range 0 to 15.5). This missingness was structured rather than random: 319 of the 394 excluded samples were brain tumors, and all 280 samples annotated only as Neoplasm, Unknown, Tumor_cells or Malignant_tumor lacked an age. The 102 distinct diagnostic strings were collapsed to 74 entities by pooling spelling variants and subtype synonyms, retaining as separate entities those with distinct clinical identity. Of these, 31 were represented among the aged tumors, and group level results are reported for the 24 entities and 8 anatomical sites with at least ten tumors (1,685 and 1,720 tumors, respectively). Paired primary and relapse tumors were available for ten patients, all from TARGET and profiled on HM450; in one neuroblastoma the relapse sample was bone marrow.

### Epigenetic clocks

Horvath DNAmAge was computed from the 353 published coefficients and intercept^8^. In the adult cohort, missing beta values at clock CpGs were imputed by the CpG mean; only two normal samples (one LUAD and one KIRC) lacked more than 28 of the 353 CpGs, and excluding these two pairs left all results unchanged (Supplementary Table 6b). In the pediatric cohort DNAmAge was estimated with the methylclock R package^38^, with cell type correction disabled, after excluding unmeasured clock CpGs and samples with remaining missing values; pediatric acceleration was defined as DNAmAge minus chronological age at diagnosis and is not directly comparable to the adult tumor minus normal difference. epiTOC1, epiTOC2, HypoClock and stemTOC were computed following the published implementations^11–14^, and epiTOC2 was used to estimate both the cumulative number of stem cell divisions and, with chronological age, the intrinsic division rate per year^12^. Per sample CpG coverage of each clock is given in Supplementary Table 31. Because coverage differs between platforms, pediatric mitotic clocks were recomputed on clock CpGs present in at least 95% of samples on all three arrays (epiTOC1 252 of 385, epiTOC2 114 of 163, HypoClock 249 of 678 and stemTOC 217 of 371 CpGs; ρ with full set estimates 0.94 to 0.99), and all pediatric mitotic comparisons were made within platform.

### Tumor purity and cell type composition

Epithelial, fibroblast and immune fractions were estimated for every adult sample by robust partial correlation with EpiDISH^27^, using the centEpiFibIC.m reference for all cancer types and centEpiFibFatIC.m for BRCA. Tumor purity was taken from the consensus purity estimate (CPE)^22^, available for 490 of 506 tumors; ESCA and four BRCA tumors lack a CPE. EpiDISH epithelial fractions were validated against CPE within each cancer type; in LIHC the epithelial fraction did not correlate with CPE (ρ = 0.13), and LIHC was excluded from composition adjusted models (Supplementary Table 7).

### Heterogeneity of methylation change

For each pair, CpGs with missing values in either sample were removed and the remaining CpGs were grouped into probe clusters, merging consecutive probes separated by no more than 500 bp^31^. For each cluster we computed Δβ as the tumor minus normal difference in mean beta value, positive values denoting hypermethylation in the tumor. Per patient summaries were the signed mean of Δβ, its standard deviation, the fraction of clusters with |Δβ| above 0.05, 0.1 and 0.2, and a discordance metric, defined as the share of clusters with |Δβ| above the threshold whose direction is opposite to the patient’s majority direction, ranging from 0 (all changes in one direction) to 0.5 (balanced). The number of components describing the distribution of |Δβ| was estimated with a Gaussian mixture model. For comparison with the earlier analysis, the same summaries were computed on non overlapping 5 Mb windows containing at least 20 CpGs, with windows classified as hypermethylated or hypomethylated when Δβ exceeded one standard deviation of the window Δβ distribution within the sample.

### X chromosome analysis

For each patient, the rate of changed units on the X chromosome was divided by the rate across autosomes, for all changes and separately for hypermethylation and hypomethylation, and enrichment was stratified by donor sex. X copy number and loss of heterozygosity in each primary tumor were obtained from open access ASCAT allele specific segments^39^ for 527 of 531 patients. Male X copy number estimates were implausible (median relative copy number 0.23) and were not used. Female tumors were stratified by X loss of heterozygosity and restricted to X copy number neutral tumors as a sensitivity analysis.

### Proliferation, genomic and clinical data

GDC STAR gene counts for the primary tumor and solid tissue normal samples of the 506 pairs were converted to log2 transcripts per million for protein coding genes, yielding tumor data for 503 patients and tumor and normal data for 273. The primary proliferation score was the mean log2 transcripts per million of the nine gene mitotic index of Yang et al. (CDKN3, ILF2, KDELR2, RFC4, TOP2A, MCM3, KPNA2, CKS2 and CDK1)^11^; rank based single sample scores of the Hallmark G2M checkpoint and E2F target gene sets^28^ and the mean within type z score of MKI67, PCNA and MCM2 to MCM7 served as sensitivity measures. Ploidy and whole genome doubling were taken from PanCanAtlas ABSOLUTE calls^40^ (489 of 506 tumors) and the aneuploidy score from Taylor et al.^41^ (488 tumors). Telomere content was taken from Barthel et al.^29^ as the log2 ratio of tumor to solid tissue normal estimates from the same library type, preferring whole genome over low pass over exome sequencing, without use of blood normals (361 pairs). Grade and stage were obtained from the TCGA Pan-Cancer Clinical Data Resource^42^, with GDC records as fallback, and analysed where at least two levels contained eight or more patients. Four pairs flagged by PanCanAtlas sample quality control were retained (Supplementary Table 16).

### Statistics and reproducibility

Tumor minus normal clock differences were tested per cancer type with two sided paired Wilcoxon signed rank tests, with Benjamini and Hochberg correction across cancer types for each clock^43^. Correlations are Spearman ρ unless stated otherwise; pooled correlations were computed within cancer type. Composition adjusted clock differences were estimated by regressing the tumor minus normal clock difference on the tumor minus normal fibroblast and immune fractions (and adipocyte fraction in BRCA) with heteroscedasticity consistent (HC3) standard errors, so that the intercept estimates the clock difference at equal composition; as a second sensitivity analysis, tests were repeated in tumors with a CPE of at least 0.6. The association between clocks and heterogeneity was tested in 490 pairs from 12 cancer types with a CPE by regressing each heterogeneity metric on the standardised clock difference, signed mean Δβ, absolute mean Δβ, log standard deviation of Δβ, CPE, age, sex and cancer type, fitted as a linear mixed model with a random intercept for cancer type, or with cancer type as a fixed effect where the mixed model did not converge; P values were corrected across all 25 tests. Heterogeneity metrics were related to purity and composition in a pooled model with cancer type fixed effects and HC3 standard errors. Tumor minus normal clock differences were regressed on tumor proliferation with CPE, changes in epithelial, fibroblast and immune fractions, signed mean Δβ, age, sex and cancer type as covariates, with continuous variables standardised within cancer type, HC3 standard errors and Benjamini and Hochberg correction; LIHC composition terms were set to zero. These models were repeated within cancer type, with the G2M score, with the tumor minus normal proliferation difference, in tumors with a CPE of at least 0.6, without the signed mean Δβ, without CPE including ESCA, excluding KIRC and excluding the two low coverage pairs. Variance was partitioned by drop one partial R² for blocks of proliferation, composition (CPE and the three fraction changes), signed mean Δβ, genomic features (aneuploidy score and whole genome doubling) and, in pairs with telomere data, telomere ratio with library type. Incremental information was tested by likelihood ratio tests adding the standardised clock difference to ordinal logistic (KIRC grade), logistic (late stage, pooled over KIRC, BRCA, KIRP and THCA; whole genome doubling) or linear (aneuploidy score, excluding THCA and PRAD) models containing proliferation, composition, signed mean Δβ, age, sex and, in pooled models, cancer type; grade in HNSC and LIHC was not reported because of separation. X enrichment was tested with paired Wilcoxon tests of X against autosomal rates. In the pediatric cohort, Horvath acceleration was tested against zero with the Wilcoxon signed rank test and compared between entities and sites with the Kruskal Wallis test; mitotic clocks were compared between entities and related to age within each entity and platform with at least ten tumors, with correction across 112 tests, and primary and relapse tumors were compared with exact paired Wilcoxon tests. Adult analyses were performed in Python 3.7.0 and pediatric Horvath analyses in R (v4.4.0).

## Data availability

The source data are available through the GDC portal from the TCGA, CGCI, TARGET and HCMI projects. ABSOLUTE ploidy, aneuploidy scores and clinical annotations are available from the PanCanAtlas publication resources^40–42^, and telomere content estimates from Barthel et al.^29^. Source data for plots are available in the supplementary files.

## Code availability

All code is provided in a repository (https://github.com/Subhayan18/TCGA_Epigenetic_age).

## Ethics statement

This study used publicly available nucleic acid sequencing data obtained through the GDC portal. No new samples were collected, and no experimental procedures involving human participants were performed. All data were fully anonymised prior to access, and no personally identifiable information was available to the researchers.

## Acknowledgements

Computational resources were provided by the National Academic Infrastructure for Supercomputing in Sweden (NAISS), funded by the Swedish Research Council. Data storage and analysis were carried out using the COSMOS secure computing infrastructure provided by LUNARC. This study was funded by a Nilsson Ehle Foundation grant from the Royal Physiographic Society of Lund, Sweden.

## Author contributions

S.C. conceived, acquired resources and supervised the study. A.M. analysed and interpreted the data and maintains the code repository. Both authors wrote and reviewed the manuscript.

## Generative AI disclosure

The authors declare use of generative AI in editing manuscript text and generating code documentation.

