## Supplementary Figure 1-3 for "Replicative and chronological epigenetic age track genome instability in opposite directions across cancers"

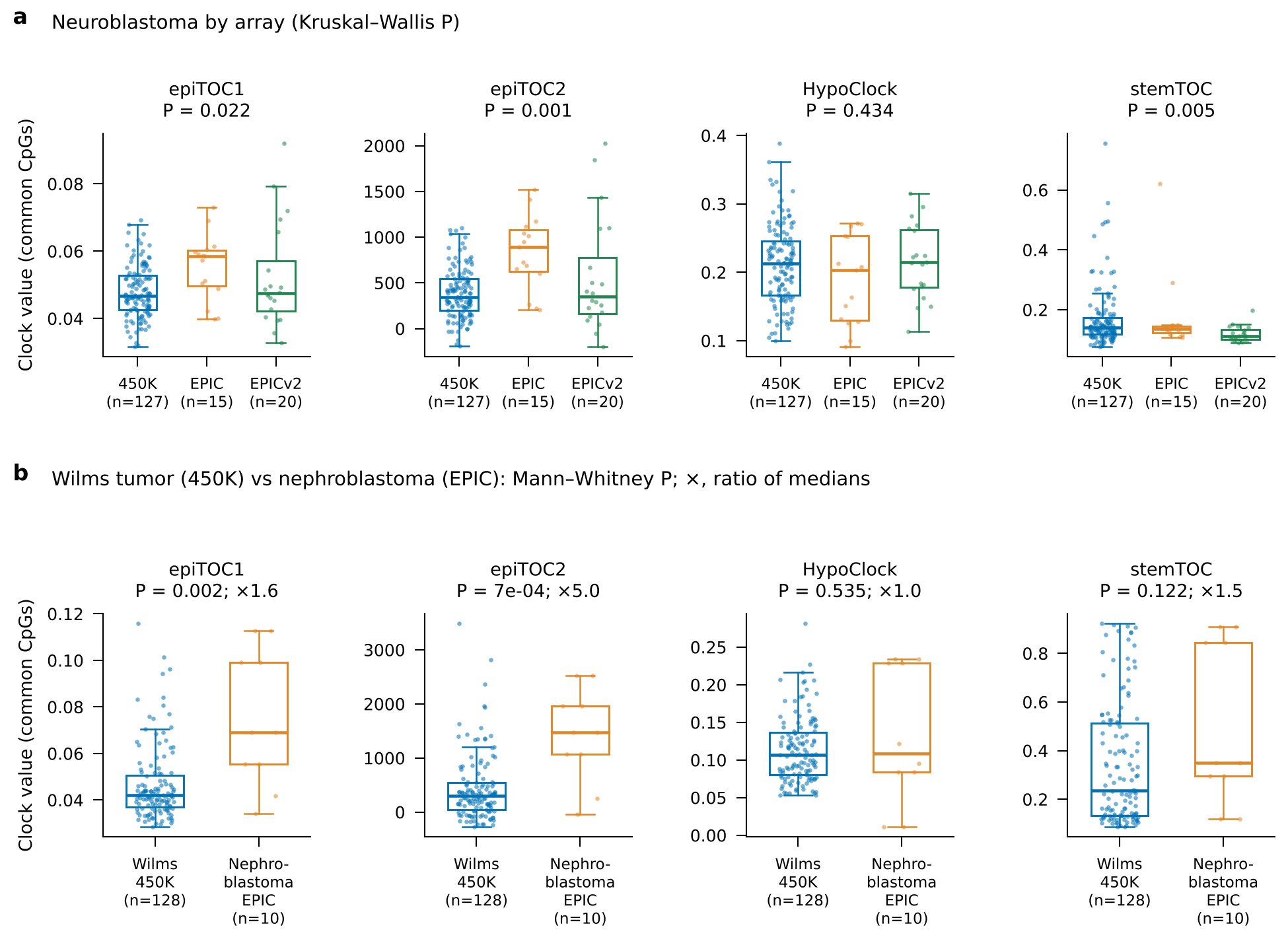

**Supplementary Fig. 1 | Platform effects on pediatric mitotic clocks.** a, Neuroblastoma clocks (common CpGs) on HM450 (n = 127), EPIC (n = 15) and EPICv2 (n = 20); Kruskal–Wallis P. b, Wilms tumors on HM450 (n = 128) versus nephroblastomas on EPIC (n = 10); Mann–Whitney P, and ratio of medians (×). Boxes show median and interquartile range; points are samples.

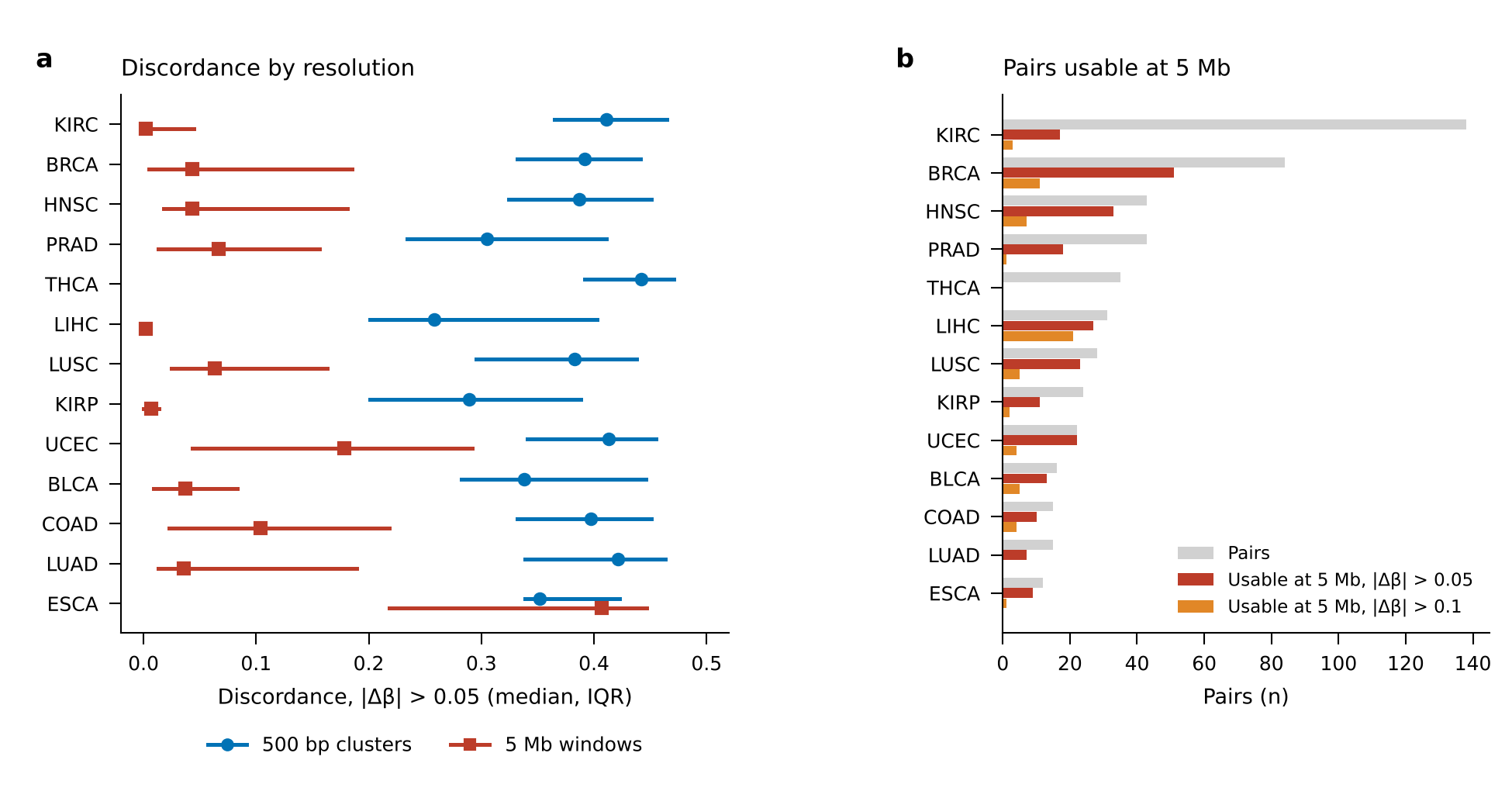

**Supplementary Fig. 2 | Discordance of methylation change at 500 bp and 5 Mb resolution.** a, Median and interquartile range of per-pair discordance (|Δβ| > 0.05) by cancer type; 5 Mb values are shown for types with usable pairs. b, Total pairs and pairs usable at 5 Mb (≥ 50 changed units). Source: Supplementary Table 24.

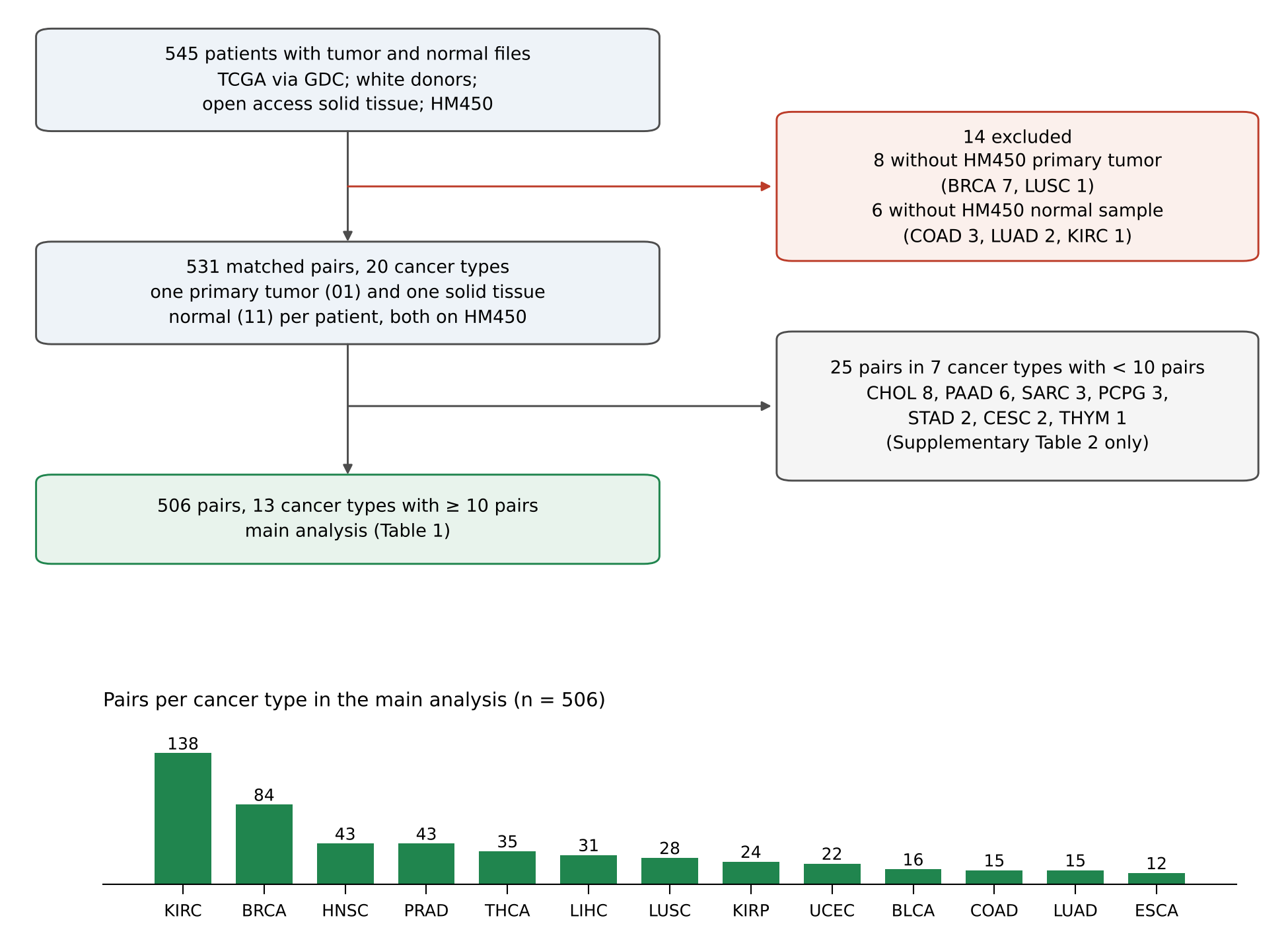

**Supplementary Fig. 3 | Construction of the adult matched tumor–normal cohort.** 545 patients with tumor and normal files, 14 excluded for lacking an HM450 primary tumor (8) or normal sample (6), giving 531 pairs in 20 cancer types; 506 pairs in 13 types with ≥ 10 pairs formed the main analysis. Bars show pairs per type. Source: Supplementary Tables 1 and 2.
